# Adaptive pangenomic remodeling in the *Azolla* cyanobiont amid a transient microbiome

**DOI:** 10.1101/2025.03.24.644982

**Authors:** David W. Armitage, Alexandro G. Alonso-Sánchez, Samantha R. Coy, Zhuli Cheng, Arno Hagenbeek, Karla P. López-Martínez, Yong Heng Phua, Alden R. Sears

## Abstract

Plants fix nitrogen in concert with diverse microbial symbionts, often recruiting them from the surrounding environment each generation. Vertical transmission of a microbial symbiont from parent to offspring can produce extreme evolutionary consequences, including metabolic codependence, genome reduction, and synchronized life cycles. One of the few examples of vertical transmission of N-fixing symbionts occurs in *Azolla* ferns, which maintain an obligate mutualism with the cyanobacterium *Trichormus azollae*—but the genomic consequences of this interaction, and whether the symbiosis involves other vertically transmitted microbial partners, are currently unknown. We generated high-coverage metagenomes across the genus *Azolla* and reconstructed metagenome assembled genomes to investigate whether a core microbiome exists within *Azolla* leaf cavities, and how the genomes of *T. azollae* diverged from their free-living relatives. Our results suggest that *T. azollae* is the only consistent symbiont across all *Azolla* accessions, and that other bacterial groups are transient or facultative associates. Pangenomic analyses of *T. azollae* indicate extreme pseudogenization and gene loss compared to free-living relatives—especially in defensive, stress-tolerance, and secondary metabolite pathways—yet the key functions of nitrogen fixation and photosynthesis remain intact. Additionally, differential codon bias and intensified (rather than relaxed) selection on photosynthesis, intracellular transport, and carbohydrate metabolism genes suggest ongoing evolution in response to the unique conditions within *Azolla* leaf cavities. These findings highlight how genome erosion and shifting selection pressures jointly drive the evolution of this unique mutualism, while broadening the taxonomic scope of genomic studies on vertically transmitted symbioses.

## INTRODUCTION

Plants partner with a wide variety of microbes to secure nutrients, tolerate stress, and regulate their development [1]. In most cases, beneficial plant symbionts are recruited by new generations from the surrounding soil or aquatic microbiota (termed horizontal transmission), while vertical transmission from parent to offspring appears rare [2]. However, the selective pressures and genomic consequences of obligate, vertically-transmitted symbioses can be extreme [3, 4], making such associations important to the study of the evolution of symbioses. The consequences of such associations include the evolution of metabolic codependence [5], life-cycle synchronization [6], and horizontal gene transfer [7]. Understanding the evolutionary pathways and fitness consequences of these modifications is crucial for efforts to control or engineer beneficial, long-term plant-microbe interactions for improving food security, carbon capture, and ecosystem restoration [8, 9].

A particularly extreme example of plant-microbe mutualism is found in the aquatic *Azolla* ferns (Salviniaceae). These ferns grow on the surfaces of still, fresh waters on every continent barring Antarctica. Despite this widespread distribution, there are seven recognized species in the genus, arranged into two taxonomic sections [10, 11]. The first section, *Azolla* sect. Azolla is presumed native to the Americas and possibly Oceania, and contains five species: *A. caroliniana, A. microphylla, A. mexicana, A. filiculoides, A. rubra*, which are now widely naturalized outside of their native ranges. The second section, *Azolla* sect. Rhizosperma includes *Azolla pinnata* and *Azolla nilotica.* The plants primarily reproduce clonally, though sexual reproduction is regularly observed in nature. As N-fixers, *Azolla* are widely used as a biofertilizer [12], a protein supplement for animals [13], and as water purifiers[14]; and have even been implicated to have played a role in global cooling during the Eocene epoch [15].

All *Azolla* species host a nitrogen-fixing endophytic cyanobacterium in hollow cavities within most of their minute leaves. This cyanobacterium—named *Trichormus azollae* Komárek & Anagnostidis 1989 (syn *Anabaena azollae, Nostoc azollae*)*—*is filamentous and heterocystous, closely resembling free-living members of the Nostocaceae [16]. Relative to most host-microbe mutualisms, the *Azolla*-*Trichormus* symbiosis is unique in its mode and efficiency of transmission. For one, *T. azollae* appears incapable of autonomous growth outside of its host and has evolved a complex, multi-stage life history that persists through both clonal (sporophytic) and sexual reproductive (gametophytic) life stages of the fern [17]. This synchronized developmental cycle, involving motile, vegetative, and dormant phases in the bacterium, results in a maternal transmission with near-perfect fidelity and hints at considerable adaptive divergence from its free-living relatives in the genera *Nostoc* and *Anabaena*. The first sequenced genome of *T. azollae* revealed extreme levels of pseudogenization, loss of genes predicted to be critical for life outside the host, and a carbohydrate transport system rarely encountered in cyanobacteria [18] and may represent the incipient stages of an endosymbiotic event that may have been similar to the evolution of the reduced N-fixing ‘nitroplast’ organelle of certain marine algae [19].

However, it is unclear whether these trends extend across the entire *T. azollae* lineage, and whether other, more cryptic evolutionary consequences — such as shifts in selection regimes on particular genes — have also occurred, as they have in many intracellular arthropod symbionts [20, 21]. Further, a suite of additional microbial taxa also resides inside the leaf chambers. Some of these bacteria have been implicated in N fixation or denitrification, but do not appear to carry out the process within the leaf cavity [22] and may themselves be vertically-transmitted symbionts of the ferns [23]. It remains unknown whether a core microbiome, beyond *T. azollae*, exists in the leaf pockets across the *Azolla* lineage and, if so, what its characteristics are.

We performed comparative metagenomic, pangenomic, and phylogenomic analyses to investigate two key questions. First, is there evidence of a shared, potentially co-diversifying *Azolla* leaf pocket microbiome across different host fern strains? Second, how do the primary *Trichormus azollae* symbionts differ from their free-living cyanobacterial relatives in terms of functional genome content, gene loss, codon bias, and natural selection regime? By generating multiple metagenome replicates for each *Azolla* species, we achieve unprecedented statistical power to identify the genes, functions, and microbial taxa essential for maintaining this unique and ecologically important mutualism.

## MATERIALS AND METHODS

### Specimen sourcing and sequencing

Living cultures of *Azolla* ferns were sourced from the *Azolla* germplasm collection kept at the Dr. Cecilia Koo Botanical Conservation Center (KBCC) in Gaoshu Township, Taiwan. This collection had been maintained since the 1980s in indoor growth chambers at the International Rice Research Institute (IRRI) in Los Baños, Philippines until its transfer to KBCC in 2017. Strains from this collection had been sourced from around the world and have been maintained as inbred lines. From this collection, we selected at least two individuals of each species that appeared healthiest (with the exception of *A. nilotica,* which was unavailable). In addition, we sourced a wild strain of what was presumed to be *Azolla filiculoides* from the University of California Botanical Garden, USA (37.875, −122.239) and *Azolla* sec. Azolla from Brazos Bend State Park in SE Texas, USA (29.379, −95.614), and *Azolla* sec. Azolla and *Azolla pinnata* growing in outdoor ponds at the USDA Invasive Plant Research Lab in SE Florida, USA (26.083, −80.242). Upon entering our lab, each strain was briefly surface sterilized with a dilute 10% bleach solution, thoroughly rinsed in sterile water, and moved into sterile plant culture containers filled with “IRRI2” nutrient medium [24]. These were allowed to grow for 4-8 months in a plant growth chamber with 16h:8h 26°/21° day/night cycle at 60% relative humidity and 150 µmol m^-2^ s^-1^ light intensity.

We isolated and concentrated leaf pockets from the host plant to enrich microbial DNA using a modified enzymatic digest method [25], followed by a standard phenol-chloroform DNA extraction on the resulting material. The steps are detailed in the **supplemental methods**. Purified leaf pocket metagenomes were sequenced on an NovaSeq 6000 (Illumina, Inc) (2×151 bp paired-end sequencing) at the US Department of Energy’s Joint Genome Institute (JGI) using their low-input metagenomic workflow [26]. Reads were quality-filtered, error-corrected, assembled using metaSPAdes [27], and mapped to assembled contigs using BBMap [28]. Feature prediction and functional annotation was carried out using PROKKA v.1.14 [29] and eggNOG mapper v.2 [30], respectively, after which predicted genes were clustered into orthologous groups using orthofinder software. Finally, contigs were binned into metagenome-assembled genomes (MAGs) using MetaBat2 v.2.15 [31] and checked for completeness with CheckM v.1.1.3 [32]. Only MAGs surpassing the ‘medium quality’ criterion of metagenome-assembled genomes (MAGs) were retained (> 50% completion, < 10% contamination) [33]. Raw and processed data are hosted on the Joint Genome Institute Genome Portal under proposal number 503794 (genome.jgi.doe.gov).

### Metagenome processing & analysis

Phylogenetic assignment of the MAGs was carried out using the PhyloPhlAn 3.0 [34] pipeline, which identifies and aligns shared markers from unknown MAGs with those from a large database of over 150,000 MAGs and 80,000 reference genomes. These alignments are used to taxonomically classify the MAGs and identify and align common marker genes, which were concatenated into amino acid alignments for partitioned phylogenetic reconstruction. We used the related pipeline MetaPhlAn4 [35] to investigate overall patterns of diversity among all contigs in each metagenome. This pipeline attempts to classify metagenomic reads based on a reference-database of ∼5.1 million clade-specific marker genes and outputs a taxonomic abundance profile. As diversity was very low for all metagenomes and sequencing depths were comparable, rarefaction of metagenomic reads was not carried out, and there was no relationship between sequencing depth and subsequent derived diversity metrics.

Two *Azolla* host plant phylogenies were estimated based on genomic SNPs and plastid marker sequences. Metagenomic reads were aligned to either the entire *Azolla filiculoides* reference genome [36] or six plastid markers (*rbcL, atpB, rps4, trnL-trnF, trnG-trnR,* and *rps4-trnS*) (one or more for every *Azolla* species) [11] using bowtie2 v.2.5.2 [37] with the --sensitive and --local options enabled. For each mapping, variant calling was carried out using mpileup in samtools v.1.21 [38]. For the full genome mapping, variants of low quality (<30) or those classified as multiallelic, monomorphic, or within 10 bases of another SNP were removed, after which SNP alignment was carried out using vcf-kit v.0.3 [39]. Reads mapped to the plastid references were also processed through mpileup and used to generate consensus sequences which were then aligned on a per-locus basis to the existing plastid reference alignment using the --add and --keeplength options in MAFFT v.7.520 [40], after which they were concatenated. Maximum likelihood-based phylogenetic inference was carried out using raxml-ng [41] with 10 random starting trees and nonparametric bootstrapping over 200 iterations. Substitution models were selected after preliminary likelihood ratio tests: GTR+G for plastid marker sequences, GTR+G+ASC_LEWIS for genomic SNPs, and LG+G8+F for the MAG partitioned amino acid alignment. Both *Azolla* trees were generally concordant, recovering the two monophyletic *Azolla* sections and two *caroliniana/microphylla/mexicana* and *filiculoides/rubra* complexes within sect. *Azolla*, with all inner branches having acceptable bootstrap support.

The R package phyloseq [42] was used to characterize the leaf pocket communities based on taxonomic assignments from MetaPhlAn4. The richness and Shannon diversities of samples were compared within leaf pockets of the two major *Azolla* clades. The R package rbims v.0.0.0.9 [43] was used to generate a database of functional predictions from the annotated MAGs. This R package organizes annotated sequences into a hierarchical database based on their KEGG IDs [44] and COG functional annotations [45] output by EggNOG mapper, which can then be used to compare metabolic pathways between genomes or metagenomes.

### Selection, gene loss, and codon bias across the T. azollae pangenome

Pangenomes of the *T. azollae* MAGs and their free-living relatives were generated using anvi’o v.8 [46] using the bioinformatic pipeline described in [47], wherein clusters of orthologous genes were aligned among related *T. azollae* genomes and visualized. We then calculated functional enrichment of particular gene clusters between clades of *T. azollae* and between *T. azollae* MAGs and the free-living outgroup genomes using the statistical test described in [48]. This test fits logistic regressions to gene cluster occurrences with focal group category as a predictor (e.g, *T. azollae* vs. outgroup genomes) and outputs false-discovery rate-corrected significance values.

Cospeciation between *Trichormus azollae* and *Azolla* was assessed using the Procrustean Approach to Cophylogeny (PACo) method, which has been shown to outperform related methods in minimizing types I and II error rates [49]. The PACo approach compares the residuals of two phylogenetic distance matrices subjected to Procrustes analysis with the residuals of a randomly permuted association matrix. Our choice of permutation algorithm (“r0”) assumed that the evolution of *T. azollae* tracked the evolution of the host plant. This permutation was repeated 1000 times returning a *p-*value for the null hypothesis that the observed cophylogenetic signal is no different than chance alone.

We investigated pseudogenization of coding regions in all MAGs of our metagenome, focusing specifically on those of *Trichormus azollae.* We used all high-quality MAGs as well as additional cyanobacterial genomes including *T. azollae* GCF_000196515.1 [18], *Nostoc punctiforme* GCF_000020025.1 [50], *Nostoc* PCC7120 GCF_000009705.1 [51], *Nodularia spumigena* GCF_003054475.1, and *Trichormus variabilis* GCF_009856605.1, of which all except *T. azollae* are free-living, heterocystous cyanobacteria isolated from aquatic habitats or plant surfaces. MAGs and reference genomes were processed through Prokka [29] to identify genes and CDs, which were then run through the Pseudofinder v.1.1 pipeline [52] using the default --annotate option against a reference database containing all cyanobacterial protein sequences available on the UniProt server (accessed on 08/14/22) [53]. We attempted to predict the original function of these pseudogenes via BLAST search against a database of all cyanobacterial proteins and then annotating the top hits from this database. This approach permits the annotation of pseudogenes that have lost crucial functional sites, making it otherwise challenging to predict their original functions. Counts of orthologous genes and pseudogenes in each cyanobacterial genome were modeled using negative binomial generalized linear models for comparison across taxonomic clade, COG categories, and pangenomic core-accessory regions.

We also quantified codon bias in orthologous genes using the ‘mean expression level predictor’ (MELP) metric with the coRdon R package [54], which measures codon bias in intact genes relative to those of known highly expressed genes (here, ribosomal proteins) and positively correlates with transcript abundances [55–57]. MELP values above 1 indicate codon frequencies match those of ribosomal genes while values below 1 indicate a closer match to genome average codon frequencies. Intact genes were further analyzed for signatures of selection. To do this, we first clustered sequences from intact genes into orthologous groups using Orthofinder v.2.5.5 [58]. Amino acid sequence alignments of each orthogroup were created with MAFFT and used for gene tree estimation with raxml-ng. Codon-aware nucleotide alignments were then generated using pal2nal software [59]. Tests for episodic positive (diversifying) selection on *T. azollae* genes were carried out using BUSTED (Branch-Site Unrestricted Statistical Test for Episodic Diversification) [60] in the HyPhy v.2.5.62 [61] software package, with default parameters and allowing for substitution rate variation. This analysis uses a likelihood ratio test to assess whether a rate-variable codon model is a better fit to the data (a subset of test branches in an orthologous gene tree) than a ‘null’ model constrained to disallow positive selection, which is characterized by a ratio of nonsynonymous to synonymous mutations greater than 1. We then tested for differential selection regimes on orthologous genes between *T. azollae* and the free-living outgroups listed above using the BUSTED-PH approach [61]. This first involves conducting a BUSTED test on groups of gene tree branches belonging to tips with particular phenotypes (here, *Azolla* symbionts and free-living bacteria). It compares the statistical likelihood of a universal *d*N/*d*S distribution over the entire gene tree to one where the two phenotypes have different distributions using a likelihood ratio test. Orthogroups failing this differential selection test still show evidence of selection associated with the focal phenotype. This may be due to the influence of outliers in either the focal or background branches, as variation in the relatively low number of branches in our comparison can increase the probability of type II (false negative) error.

Finally, we used HyPhy’s RELAX test to assess whether either purifying or positive selection on a particular orthologous group was either relaxed or intensified in the *T. azollae* clade as a consequence of their symbiotic lifestyle [62]. This test compares an unconstrained null *d*N/*d*S model fit to that of a more complex model containing a *selection intensity parameter* (*k*) which imposes a shift in the null model between focal and background gene tree branches. Values of *k* > 1 indicate selection on the focal branches (*T. azollae*) has intensified relative to the background (free-living Nostocales), whereas 0 < *k* < 1 indicates selection has relaxed toward neutrality. Genes that were identified as having exceptionally elevated *dN/dS* from the Pseudofinder analysis were not included in this test, but are considered cryptic pseudogenes that are treated as such in our analyses [63, 64]. Only orthologues present in all free-living outgroups and at least 22 of the 24 *T. azollae* MAGs were included. For all *p*-values, false discovery rates were controlled using the Benjamini-Hochberg step-up procedure [65].

## RESULTS

### Microbial community patterns

Metagenome assembly resulted in an acceptable average N50 value across all assemblies of 31 kb, with high average coverage (>300×) and mapping ratio (>95%) across all contigs. Contig binning returned 121 medium and high-quality MAGs. Statistics for these are provided in the supplement.

Between one and fourteen MAGs were recovered per *Azolla* sample, with *T. azollae* reads recovered at high relative abundances in every metagenome (**Fig. 2A**). Beyond the cyanobacterial symbionts, there was no group of bacteria consistently found within all or even the majority of leaf pockets. The highest number of non-cyanobacterial MAGs were classified to the order Rhizobiales (syn. Hyphomicrobiales), which were found across 43% of samples. The leaf cavity samples with the highest taxonomic diversity were those that were freshly collected from natural habitats, indicating that losses of bacterial populations in leaf cavities likely occurred during long-term laboratory propagation. On average, lower taxonomic diversity (in terms of raw richness and Shannon diversity) was observed in the leaf cavities of *Azolla pinnata* (sect. Rhizosperma) relative to *Azolla* sect. Azolla (richness: *t*_19.7_ *=* 2.3, *p* < 0.05; Shannon *t*_15.7_ *=* 2.3, *p* < 0.05), though the statistical significance of this comparison disappeared when wild-collected samples were excluded from the analysis. Overall, most leaf pockets contained very low microbial diversity (mean = 5.3 ± 0.2 species) (**Fig. 2)**.

**Figure 1.**
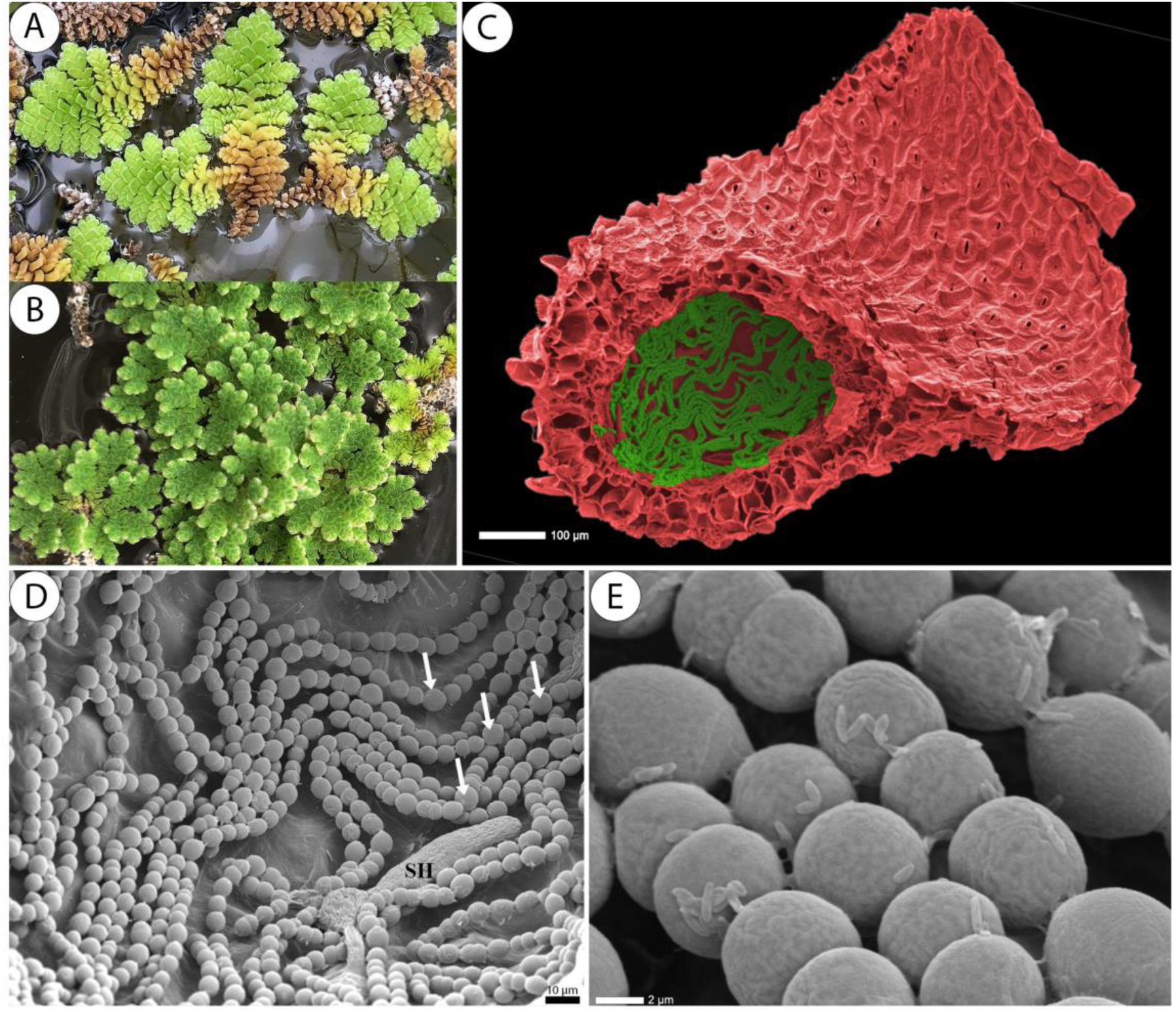
(A) Photographs of *Azolla pinnata* (sect. Rhizosperma) and (B) *Azolla caroliniana* (sect. Azolla). (C) Scanning electron micrograph of the leaf (red) and leaf pocket (green) of *A. pinnata*. (D) Close-up of *Trichormus azollae* cells within the pocket. Arrows show examples of heterocyst cells, SH is a simple hair cell. (E) Close-up of unknown bacteria attached to individual *T. azollae* cells. Details on SEM imaging are in the supplemental methods.

**Figure 2.**
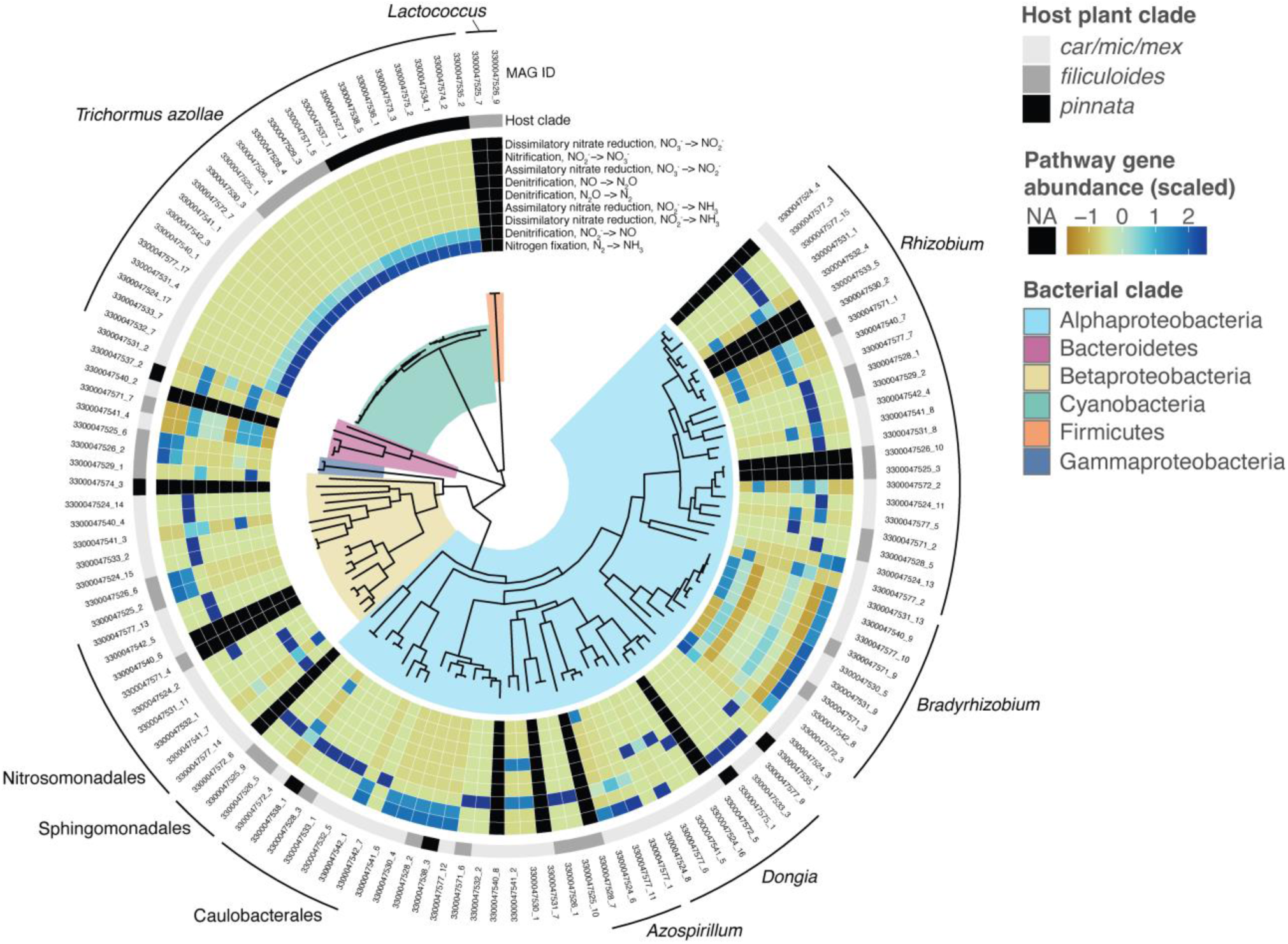
(A) Phylogenomic tree of medium and high-quality MAGs recovered from *Azolla* leaf pocket metagenomes. Branches are shaded by bacterial clade. First series of rings shows normalized KEGG ortholog gene counts within major nitrogen transformation pathways in each MAG. (B) Taxonomic profile of the leaf microbial communities detected in each leaf pocket metagenome. Note the different units for each *y-*axis. (C) Heatmap of KEGG metabolic pathways (rows) for each MAG (columns), excluding nitrogen pathways already shown in panel A. Dendrogram shows Ward’s hierarchical clustering based on metabolic similarity (including N pathways). MAG label colors follow bacterial clade colors in panel A.

The metabolic traits of the leaf pocket MAGs were quite diverse, and hierarchical clustering separated the cyanobacterial symbionts and a small number of facultative anaerobes from the majority of the MAGs (**Fig. S1**). Relative to the non-cyanobacterial MAGs, the *T. azollae* genomes were characterized by complete N-fixation pathways and an absence of chemotaxis and associated motility genes. Metabolic pathways represented in the remainder of MAGs were highly variable, and included a wide array of inorganic N transformations, including N fixation outside of *T. azollae* (**Fig. 2**). Various fixed-acid fermentative pathways were also detected, including in a homofermentative, nonmotile *Lactococcus* species. Additional putatively nonmotile MAGs included members of Chitinophagaceae detected across multiple samples. Oxidative phosphorylation pathway genes were common across most MAGs but notably reduced in those classified as *Enterobacter ludwigii, Methylovorus* sp., and the aforementioned *Lactococcus* genomes. However, there were no clear trends in metabolic traits in the set of MAGs that clearly distinguish them from other common free-living aquatic or plant-associated bacteria.

### Evolutionary genomics of T. azollae

Comparison of the *T. azollae* and *Azolla* phylogenetic trees (based on both genomic SNP and plastid markers) revealed substantial cophylogenetic signal, supporting the hypothesis that the phylogeny of *T. azollae* tracks that of its host plant (PACo SS_resid_ < 0.05, *p* < 0.0001 for both *Azolla* trees) **(Fig. 3, Fig. S3)**. Pangenomic analysis of the *Trichormus azollae* MAGs revealed a suite of 1672 conserved, intact gene clusters common to both the obligate cyanobionts and their free-living relatives, which we designated the *core pangenome* (**Fig. 3).** All single-copy genes fell into this conserved core. A second, more fragmented conserved region was also heavily shared among all genomes. Two additional clusters were also identified. The first contained the 2333 clusters exclusive to the free-living taxa while the second contained 1069 ‘accessory’ gene clusters that were inconsistently shared among *T. azollae* lineages. The gene clusters found within these core and accessory bins differed in their functional annotations, with only COG category X (mobilome/prophages/transposons) having a greater total abundance in the accessory genome than the core genome **(Fig. S2A)**. Likewise, the fractional representation of each COG category in intact clusters also significantly differed between core and accessory genomes, with gene clusters belonging to COG categories X (mobile elements), Q (secondary metabolite synthesis), V (defense), K (transcription), P (inorganic ion transport/metabolism), and T (signal transduction) being found at higher frequencies (relative to other COG categories) in the accessory set compared to the core pangenome **(Fig. S2B)**.

**Figure 3.**
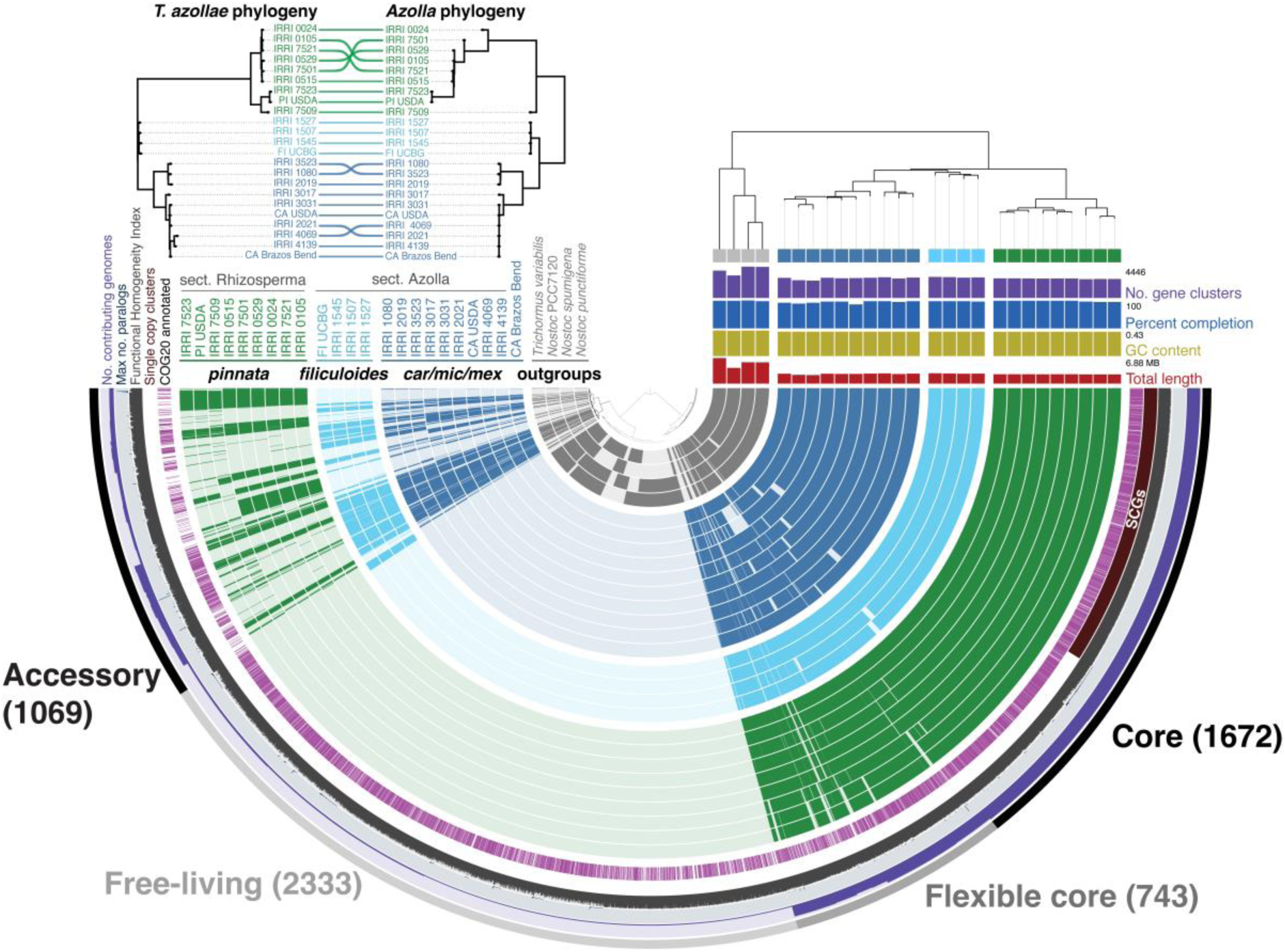
Pangenome of *Trichormus azollae* symbionts and a selection of closely-related free-living cyanobacterial genomes. Inset graph shows phylogenomic trees of *T. azollae* MAGs and their host ferns with lines connecting the host and symbiont tips. Vertical ordering is based on phylogenetic relatedness. Inner rows on the radial display show intact gene clusters of each genome. The ‘core genome’ is the region on the right comprising 1672 gene clusters shared across all genomes. “SCGs” bar indicates the region of single-copy genes, all clustered within the core genome. The functional homogeneity index identifies relative similarity of each gene cluster with respect to the amino acid composition of each cluster among genomes.

Functional enrichment analysis of pangenomic gene clusters identified 693 (35% of the total count) KEGG-assigned gene clusters and 974 (40% of the total count) COG-annotated clusters to be statistically associated with either *T. azollae* or their free-living relatives (**Fig. 4A**). *Trichormus azollae* genomes were significantly enriched with genes regulating fimbral expression and sensing of histidine kinase — important components of attachment, biofilm formation, and secretion. Conversely, the free-living outgroup genomes were enriched with clusters coding for phototaxis, light capture, UV protection, gas vesicle formation, and osmoregulation. A few of these genes were specifically associated with individual *Azolla* lineages. Notably, two distinct fimbrial regulatory genes, *fimE* and *fimB*, were uniquely found in the sections Azolla and Rhizosperma, respectively, but were absent in the free-living outgroups. A COG-category-wise analysis of differentially enriched genes suggests that the three most enriched categories in the outgroups relative to *T. azollae* were associated with categories X (mobilome), Q (secondary metabolites), and V (defense mechanisms) (**Fig. 4B**).

**Figure 4.**
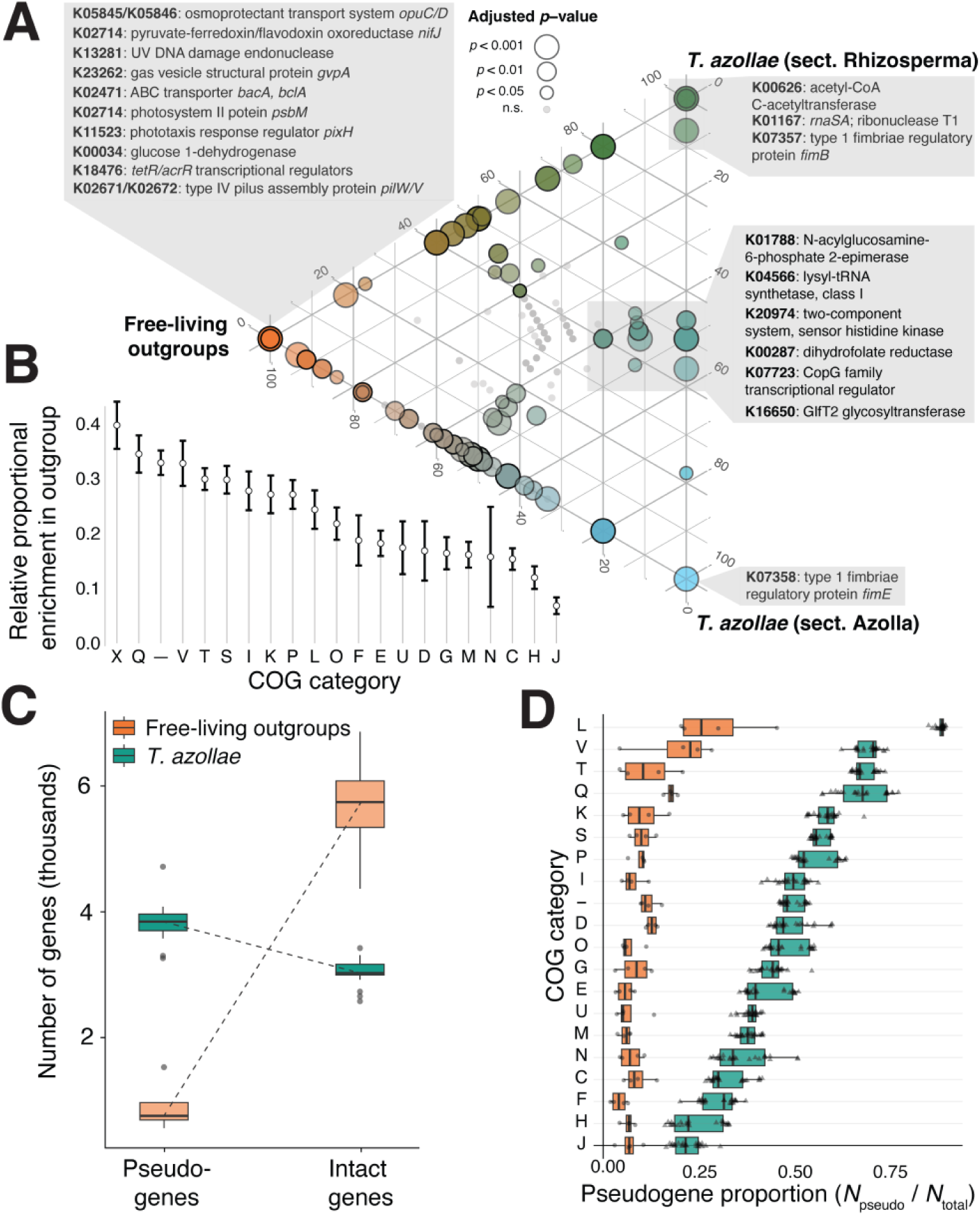
(A) Functional enrichment of gene clusters between the three focal groups. Placement of points and their coloring represent relative representation of each gene cluster among the three genomic groups. KEGG annotations of certain differentially-enriched gene clusters are shown in gray boxes. (B) Differential functional enrichment of genes by COG category. Higher values indicate fewer genes within a category were present across *T. azollae* genomes relative to outgroup genomes. (C) Comparison of pseudo- and intact gene counts between the *T. azollae* and its free-living relatives. (C) Percentage of pseudogenes in each genome as a function of COG category. Category key: L = replication and repair, V = defense mechanisms, T = signal transduction, Q = secondary structure, K = transcription, S = unknown function, P = inorganic ion transport and metabolism, I = lipid metabolism, D = cell cycle control and mitosis, O = post-translational modification and protein turnover, G = carbohydrate metabolism, E = amino acid metabolism, U = intracellular trafficking and secretion, M = cell wall/membrane/envelope biogenesis, N = cell motility, C = energy production and conversion, F = nucleotide metabolism, H = coenzyme metabolism, J = translation, X = mobilome/prophages/transposons, “–“ = no description).

Pseudogene counts were significantly higher in *T*. *azollae* compared to their free-living relatives (Neg. Binomial GLM; *Z =* 17.72, *p* < 0.0001) **(Fig. 4C)**, and, in *T. azollae,* were significantly greater in abundance than even intact genes (Neg. Binomial GLM; *Z =* - 12.79, *p* < 0.0001). The relative fractions of pseudogenes among COG categories varied significantly for both the cyanobiont and free-living outgroup genomes **(Fig. 4D)**. However, this variability was much higher for *T. azollae*, which had significantly more pseudogenes than intact genes for COG categories L (replication and repair), V (defense mechanisms), T (signal transduction), Q (secondary structure), and K (transcription), with the largest differences between *T. azollae* and outgroup genomes in replication/repair and signal transduction categories. In contrast, MAGs classified to the order Rhizobiales (the second hypothesized symbiont of *Azolla*) did not show evidence for greater fractional pseudogene representation in their genomes (*t*-test *t*_92_ = 0.8, *p =* 0.43) and had higher average intact gene counts than most other non-cyanobacterial MAGs recovered from the samples (Neg. Binomial GLM *Z =* −3, *p* < 0.005).

The mean expression level predictor (MELP) metric found 90 orthologous groups to be differentially biased (and therefore possibly differentially expressed) between *T. azollae* and free-living relatives (**Fig. 5A**). Of these, 32 had significantly higher MELP values in *T. azollae* compared to the outgroup genomes, whereas 58 were significantly lower than the outgroup. Genes with MELP scores greater than 1 in the outgroup and less than 1 in the cyanobionts include those coding for glycosidases (cd, ma nplT), cobalamin synthesis (*cobW*), and Fe-S cluster assembly proteins (*sufB*). Genes enriched in the cyanobionts relative to the outgroups include photosystem II proteins (*psbK*), phycoerythrocyanin linker proteins (*pecC*) and a putative virulence factor (*mviM*). MELP scores of orthologous genes varied between *T. azollae* and the free-living relatives in a category-dependent fashion (**Fig. 5B**). Most markedly, orthologues in the COG category U corresponding to intracellular trafficking, secretion, and vesicular transport had the greatest differences between the two groups and was the only category with significantly higher predicted expression in *T. azollae* genomes relative to the outgroup. All other statistically significant MELP comparisons favored the free-living outgroups.

**Figure 5.**
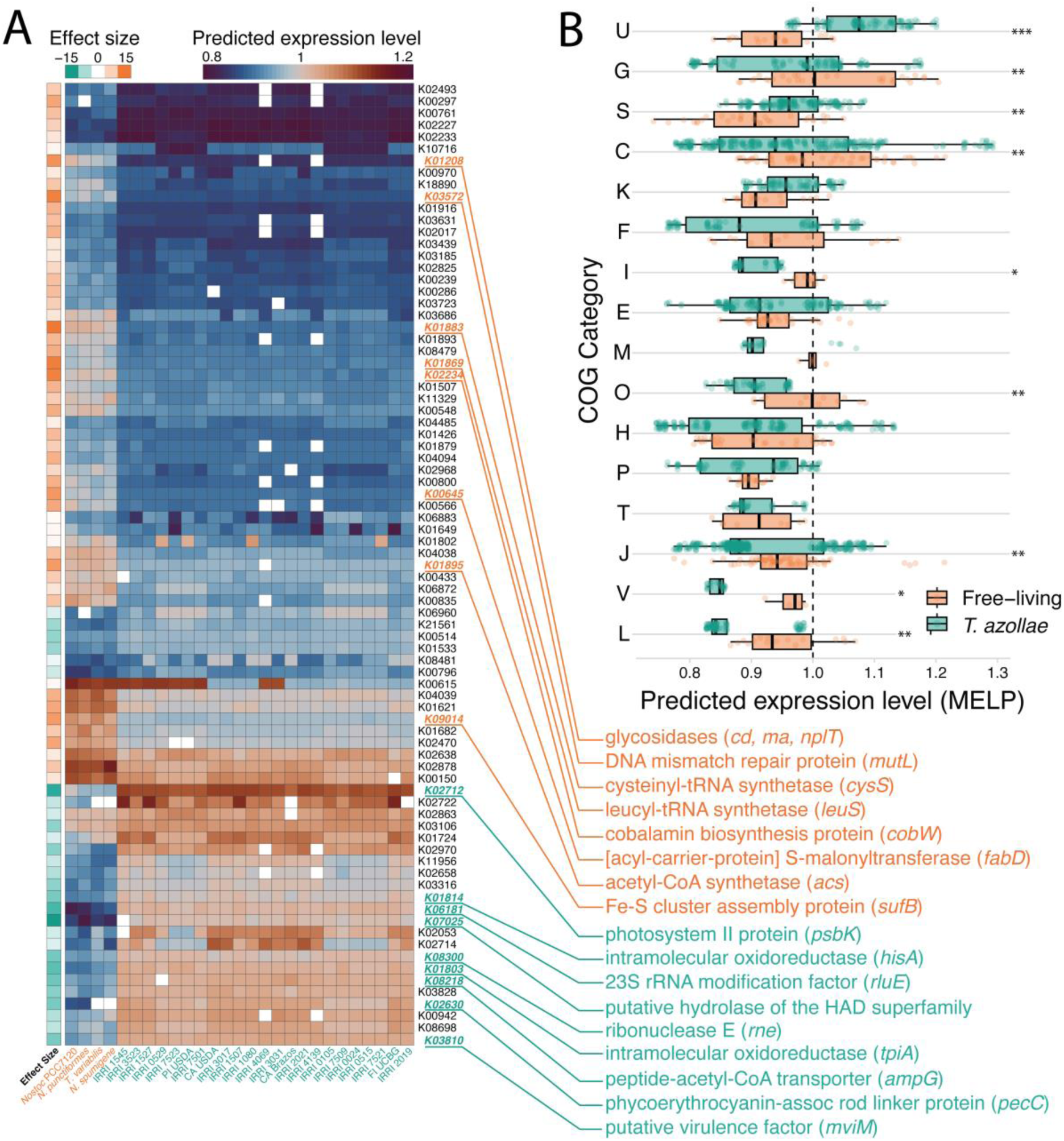
(A) Heatmap of predicted gene expression based on the MELP metric. Values greater than 1 indicate codon bias is closer to highly expressed ribosomal reference genes than genomic average codon frequencies. Rows are genes shared among *T. azollae* and free-living relatives which could be assigned to KEGG orthologs. The subset of genes shown here correspond to those with significantly different MELP scores between the two groups with substantial effect sizes (|*D*| > 0.8) indicated in the leftmost column, where negative values indicate higher MELP scores for *T. azollae* and positive values indicate higher scores for the free-living relatives. The top 10% of these genes with regard to effect size are highlighted alongside their KEGG annotations. (B). Genes with MELP scores that significantly differed between *T. azollae* and free-living genomes, ordered by COG category. Asterisks next to each line denote adjusted significance of difference among groups (*p* < *0.05, **0.001, ***0.0001). Here, for example, genes in the “U” category (intracellular trafficking and secretion) have significantly higher MELP scores than in free-living relatives. COG category definitions are listed in the Figure 4 caption.

Tests for changes in the strength of selection for orthologous genes using the RELAX procedure returned 139 orthologues (out of 3520 total) exhibiting statistically significant evidence of intensified selection (*k >* 1) and 22 showing evidence of relaxed selection (*k* < 1) (**Fig. 6A**). Genes in the test group identified as having relaxed in selection strength include many in the COG categories T (signal transduction) and L (replication, recombination and repair) and include subunit 1 of the cytochrome oxidase enzyme (*coxA/ctaD*), and two DNA replication and repair proteins (*recF* and *alkA*). Conversely, genes experiencing intensification of selection were detected across all COG categories and included circadian clock proteins (*kaiB/kaiC*), genes involved in photosynthesis (*ftrC, chlB, psbC*), transport & adhesion (*psbC, pilC, exbD*) and heterocyst pattern formation (*patA*).

**Figure 6.**
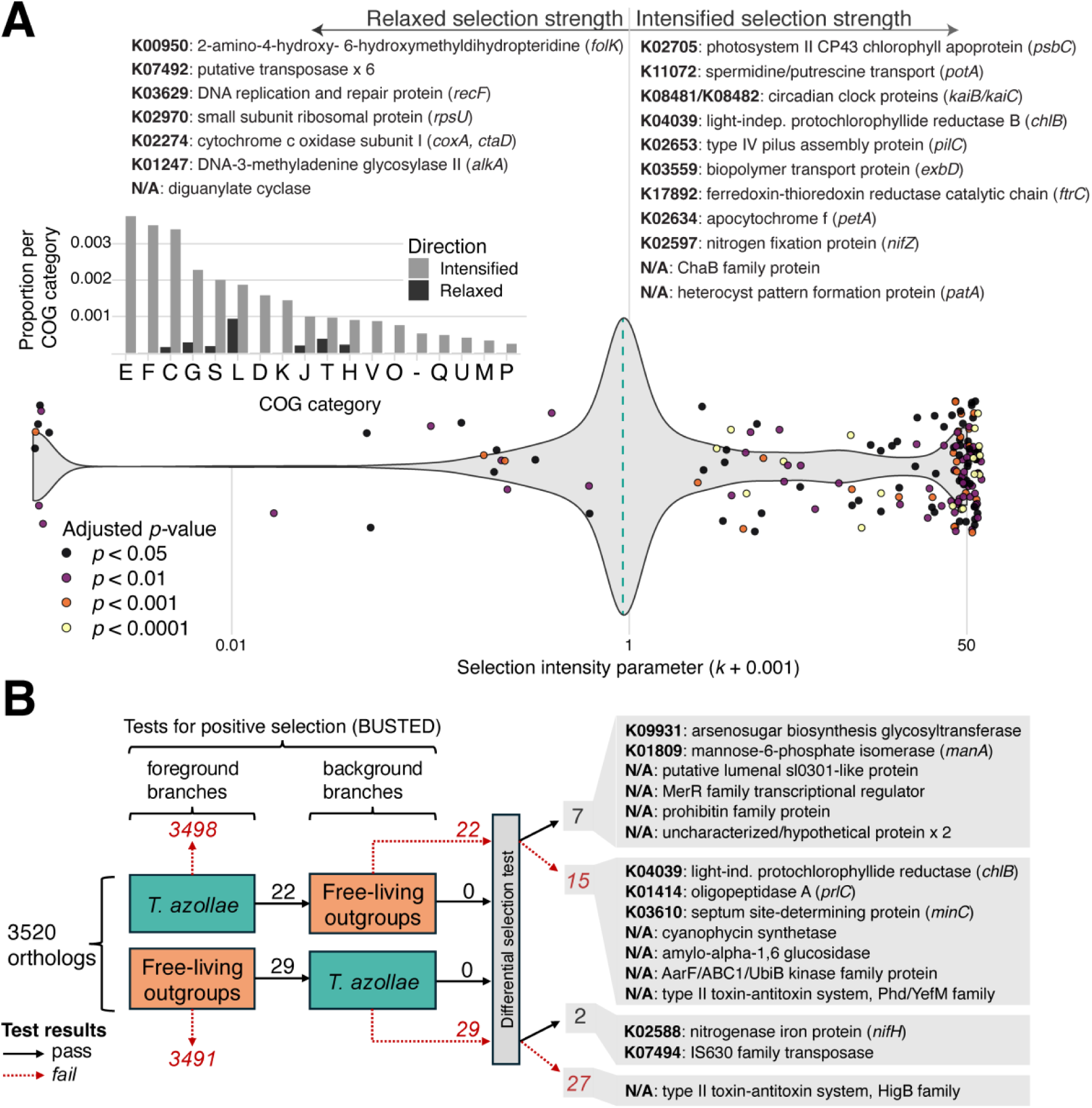
(A) Results of RELAX analysis for intensification or relaxation of natural selection on test branches (corresponding to *T. azollae* MAGs) of ortholog gene (OG) trees. The violin plot shows the overall distribution of selection intensity parameters inferred for each OG. Values greater than one indicate selection on the OG is intensified in test branches and values less than one indicate selection has relaxed. Dashed line denotes median *k* across all genes and points identify the OGs where *k* ≠ 1. Lists above the graph identify a subset of annotated OGs. Inset bar graph shows the relative distribution of OGs with relaxed or intensified selection relative to the total number of pangenomic orthogroups in each COG category. (B) Flowchart illustrating the results of the BUSTED-PH test for differential positive selection on OGs. Arrows denote the number OGs either passing or failing each statistical test for positive selection on foreground/background branches and differential selection between the two. The boxes on the right highlight the annotations of some of the OGs found to differ in selection regimes between *T. azollae* and the free-living outgroup genomes.

Subjecting these orthologues to a more specific test for positive selection under the BUSTED-PH procedure revealed a suite of 22 and 29 mutually exclusive orthologues identified as having at least one branch experiencing statistically significant positive selection in *T. azollae* and free-living relatives, respectively (**Fig. 6B**). Of these orthologues, only 7 (for *T. azollae*) and 2 (for the outgroup) showed evidence for statistically greater positive selection than the background group. Genes differentially under increased positive selection in *T. azollae* include an arsenosugar glycosyltranferase and mannose-6-phosphate isomerase, while additional genes under positive selection in this group but failing the final differential selection test include proteins playing important rolls in chlorophyll production in low-light environments (*chlB*), inhibition and directionality of cell division (*minD*), and long-term nitrogen storage (cyanophycin synthetase). In contrast, genes under significantly increased positive selection in the free-living outgroup include nitrogenase iron protein (*nifH*) and an IS630 family transposase.

## DISCUSSION

In this study, we used metagenomic and pangenomic analyses to examine the taxonomic and metabolic diversity of the symbiotic leaf pocket microbial community across much of the *Azolla* fern lineage. We focused on comparing the pangenomes of the dominant cyanobacterial symbiont, *Trichormus azollae*, with those of closely related free-living species to identify patterns in functional gene enrichment, pseudogenization, and differences in codon bias and selection regimes on specific orthologous genes.

At the community level, we encountered only one bacterial taxon — *T. azollae* (Nostocales) — that was present in leaf pockets across all *Azolla* strains, regardless of whether the plants had been recently collected from the wild or were cultured as germplasm for the past 30 years. This extreme host fidelity is due to the unique vertical transmission mechanism of *T. azollae*, which persists even in benign germplasm culture conditions due to its fitness benefits to the host [66]. While other microbial taxa were found across multiple host fern strains — most notably *Bradyrhizobium* and *Rhizobium* species — none showed a cophylogenetic signal comparable to that between *T. azollae* and their host plants. This suggests that although the conditions in the leaf pocket are suitable for the invasion and possible vertical transmission of bacterial clades with a reasonably wide range of metabolic traits, the associations of most non-cyanobacterial clades are likely transient and nonessential for host performance.

The relatively large representation of *Rhizobium* and *Bradyrhizobium* MAGs in the leaf pocket metagenomes aligns closely with the results of other recent surveys. Dijkhuizen *et al.* [22] surveyed the compositions of eight *Azolla* accessions including one of the same strains analyzed here (IRRI 3017). Here, the authors were able to recover ribosomal RNA biomarkers for the Rhizobia at comparable relative abundances to those recovered here. In agreement with our study, the authors further speculated that these taxa did not contribute to N-fixation in the leaves as they lacked the requisite genes in their MAG assemblies. Another recent study of 112 *Azolla* leaf pocket microbiomes in wild strains from California, USA (primarily members of sect. *Azolla*) also encountered members of Rhizobiales across their samples [23]. Their analysis revealed that these non-cyanobacterial members show a weaker significant cophylogenetic signal with the host fern than that of its primary cyanobiont.

The causes of this pattern, however — whether via vertical transmission or via repeated invasion of leaf pockets from distinct local source populations — remain unclear. Rhizobia are already known to epibiotically colonize the heterocysts of both free-living *Anabaena* species [67] and *T. azollae* [68]. However, none of the non-cyanobacterial MAGs showed evidence of cophylogeny with the host plant, and most possessed pathways for flagellar assembly and chemotaxis, suggesting that invasion of a leaf pocket and transient persistence by motile, free-living bacteria from the surrounding environment happens with some regularity. Invasion most likely occurs in the early stages of leaf development, when the young leaf’s pocket is briefly open to the outside environment [17]. To better clarify the mode of microbiome transmission and leaf invasion, a fluorescence *in situ* approach could be used to track the spatial location and persistence of a bacterial clade such as *Rhizobium* across life stages.

Comparison of the *T. azollae* pangenome to the genomes of close, free-living relatives revealed differences in pseudogenization, gene set enrichment, codon bias, and signatures of selection which help clarify the evolution of this cyanobacterium’s unique lifestyle. Our results confirm at the clade level the substantial pseudogenization (56%) in the *T. azollae* genome first described by Ran *et al.* [18]. The pseudogenic content of most bacterial genomes appear to range between 1% and 25% [69, 70], though our approach to pseudogene detection uses an expanded suite of criteria (most notably genes with *dN/dS* > 0.3) to detect pseudogenes [63], and therefore returns pseudogenization estimates between 10% and 30% higher than earlier methods [52]. It is hypothesized that highly-pseudogenized genomes of symbiotic bacteria are expected in recently evolved symbioses which have not been stable long enough to experience reductive genome evolution. This is likely the case in *Azolla*, as the fossil-calibrated molecular clock places divergence of the genus at 89 Ma in the late Cretaceous [11]. In contrast, highly eroded intracellular symbiont genomes such as *Buchnera* have divergence estimates of at least 150 Ma or earlier [71], though this reduction process may have already begun in *T. azollae* as the average genome size has decreased to ca. 5 Mb from 6-7 Mb in their free-living relatives [18].

Our functional annotation of these pseudogenes both confirms and expands the scope of previous genomic analyses of *T. azollae.* In agreement with Ran *et al*. [18], the highest proportions of pseudogenes were from replication, recombination, and repair (COG category L), signal transduction (T), and secondary metabolite biosynthesis, transport, and catabolism (Q) categories. Notably, the former two categories also contained the most genes experiencing relaxed selection. The relative degree of pseudogenization in signal transduction genes was especially pronounced. Genes in this category have also been disproportionately purged from numerous obligate symbionts of arthropods (e.g., *Serratia*, *Buchnera, Pantoea, Rickettsia*) [72–74]. However, this trend is less apparent in the vertically-transmitted bacteria present in the leaf nodules of angiosperms such as *Psychotria* and *Ardisia* [75, 76].

Among the orthologous genes shared between free-living cyanobacteria and *T. azollae*, many exhibited differential codon bias. Notably, genes associated with intracellular trafficking, secretion, and vesicular transport (U) were the only functional category predicted to be more optimized for expression in *T. azollae* compared to the outgroup. In contrast, genes related to posttranslational modification (O), defense mechanisms (V), and replication and repair (L) were predicted to have higher expression in the outgroup than in *T. azollae*. Additionally, the functional enrichment of COG categories related to wall, membrane, and envelope biogenesis (M) and carbohydrate transport and metabolism (G) in the *T. azollae* genomes suggests a shift in selective pressure away from environmental stress tolerance and toward adhesion and extracellular transport.

We identified hundreds of genes that exhibit signs of differential enrichment, codon evolution, or natural selection between *T. azollae* clades and free-living Nostocales. Many of these genes are linked to pathways involved in nitrogen fixation, motility, stress tolerance, photosynthesis, carbon metabolism, and defense. Major differences in the properties of these orthologs between the cyanobacterial lineages are briefly highlighted below.

### Nitrogen Fixation and heterocyst development

The *Azolla-Trichormus* symbiosis relies on effective nitrogen fixation, reflected in the higher heterocyst frequency of *T. azollae* (∼20%) compared to free-living cyanobacteria (∼5-10%) [77, 78]. This suggests a greater investment in nitrogen fixation at the expense of photosynthesis and replication. In line with this observation, we found evidence for increased selection pressure on the *patA* gene — a key regulator of heterocyst formation and patterning [79]. Additionally, cyanophycin synthetase (critical for nitrogen storage and transfer among cells) and *nifZ* (essential for MoFe protein synthesis in nitrogenase) are under positive selection in *T. azollae* but not in the outgroup. The *nifH* gene, on the other hand, appears to be experiencing increased positive selection in the outgroups, despite an earlier proteomic observation of its post-translational modification in *Azolla* leaves [80]. However, whereas our analyses focused solely on gene bodies, a large 11 kb intergenic spacer between *nifK* and *nifDH* has previously been identified [81] which may modify or hasten the translation of these nitrogenase enzymes outside selection on the gene bodies themselves [82].

### Motility, chemotaxis, and stress tolerance

Genes associated with environmental stress responses were enriched in free-living outgroups. and include osmoprotectant transporters (*opuC*/*D*), UV damage endonucleases, and efflux pump regulators *tetR*/*acrR* [83]. Motility-related genes coding for type IV pilus proteins (*pilW*/*V*), a gas vesicle structural protein (*gvpA*), and phototaxis regulator (*pixH*) were also enriched in free-living genomes. In contrast, *T. azollae* showed enrichment in fimbriae-associated adhesion proteins, an important structure in plant-cyanobacterial symbiosis [84]. The fimbral regulator *fimB* was enriched in *A. pinnata* (sect. Rhizosperma), while its counterpart *fimE* was enriched in sect. Azolla species. Interestingly *fimB* exhibits optimal performance at higher temperatures than *fimE* [85], which aligns with the increased performance of *A. pinnata* at higher temperatures relative to *A. filiculoides* [86]. In addition, genes involved in DNA repair such as *recF* and *alkA* showed evidence for relaxed selection in *T. azollae*. This evidence indicates a broad evolutionary genomic reshaping in the transition from a motile existence in a periodically stressful environment to a more stable, surface-attached lifestyle in line with the natural history of *T. azollae*.

### Photosynthesis and carbon metabolism

We anticipated genes related to photosynthesis should be under relaxed selection in *T. azollae* given that its carbon appears to be provisioned by its host plant. Genes that showed the largest differences in codon bias between the test groups included antenna protein-related genes *pecA/pecC,* and photosystem II subunits *psbM, psbY, psbX,* and *psbK* —all of which were predicted to be optimized for expression in *T. azollae*. Similarly, two PSII antenna protein genes (*psbC, psbY*) and cytochrome f (*petA*) are predicted to be experiencing increased selection pressure in *T. azollae*. This differential codon optimization and increased selection on multiple essential and conserved photosystem components in *T. azollae* is surprising and begs further investigation, since it has long been understood that while the *Azolla* ferns show especially high photosynthetic rates [87], their *T. azollae* symbionts seem to exhibit markedly lower CO₂ fixation rates compared to free-living *Anabaena* strains [80]. One possible explanation for this discrepancy is that selection on photosynthetic genes in *T. azollae* may favor increased efficiency rather than absolute carbon fixation capacity. The continued production of photosynthetic pigments and enzymes suggests that maintaining a functional photosynthetic apparatus is important for the symbiosis. This may help supplement its carbon budget during periods of host dormancy, when external carbon supply is reduced. Such a mechanism seems necessary in seasonal climates when the host plants exhibit dramatic changes in pigment composition and carbon fixation rates [88]. Additionally, the light environment within *Azolla* leaf chambers may be quite different from the spectrum experienced by free-living cyanobacteria.

Little is currently known about the physiology of *T. azollae*, since they have proven resistant to culturing efforts, and previous investigations of cyanobacterial isolates from *Azolla* have, by accident, been carried out on free-living cyanobacterial contaminants rather than the primary symbiont itself [89]. To progress on this front, our genomic data can be used to develop metabolic models which can guide the development of isolation media [90]. Further, there have been promising developments in identifying differentially expressed genes in the host fern in the presence and absence of *T. azollae* [91], but similar studies of gene expression in the symbiont are needed [92], particularly across different stages of the host plant’s life cycle. Likewise, studies linking physiological status and gene or metabolite expression *in situ* are becoming more common in *Azolla* [93–95], and the divergently evolving or eroding genes identified herein can contextualize results in this developing area of research.

The *Azolla* leaf pocket is also a useful model microcosm for investigating microbial community dynamics at high spatiotemporal resolution. Fine-scale genomic and transcriptomic assays, alongside *in situ* imaging could provide direct insights into the invasion and persistence of mixed strain assemblages in these micro-scale bioreactors. Introducing new microbes to the plant at the megaspore stage has already been successfully carried out and, quite surprisingly, has resulted in the successful introduction of a free-living cyanobacterial isolate into the leaf cavity [96]. This sets up the exciting prospect of experimentally introducing novel beneficial symbionts to either improve plant productivity or study the evolution of the symbiosis by attempting to recreate its early stages.

## Conclusions

Contrary to observations of a diverse, shared leaf pocket microbiome in wild *Azolla* ferns, our metagenomic analysis of leaf pockets from strains both recently collected and from long-term cultures do not support a persistent vertically-transmitted microbiome outside of the cyanobacterium *Trichormus* (syn. *Nostoc, Anabaena*) *azollae,* the persistent microbial symbiont of its host plant. Our findings further highlight the substantial genomic differentiation of the *T. azollae* symbiont from its free-living relatives, shaped by both selection on essential symbiotic functions and the ongoing loss of non-essential genes. The evolutionary trajectory of the *Azolla-Trichormus* symbiosis appears to be one of ongoing, pairwise co-diversification, with natural selection reshaping the cyanobiont’s role in nitrogen fixation, host adhesion, and metabolite transfer, while retaining the metabolic independence that differentiates it from more ancient intracellular symbionts. By expanding our understanding of the only known vertically transmitted nitrogen-fixing mutualism, these results contribute to a growing body of knowledge on how bacterial symbioses evolve in complex settings, particularly in the ecologically and economically important N-fixing cyanobacteria.

## ACKNOWLEDGEMENTS

The authors thank the Cecilia Koo Botanic Conservation Center (KBCC), Ken-Yu Cheng (KBCC), C. Rothfels (Utah State University), H. Forbes (UC Botanical Garden), P. Madeira (USDA), E. Pokorny (USDA) for assistance with specimen sourcing.

## Statement of authorship

DWA conceived this work. DWA, ADA-S, SRC, and ARS performed data collection. DWA, ZC, AH, KPL-M and YHP performed data analysis, and DWA wrote the manuscript with input from all authors.

## Data accessibility statement

Data are publicly available on the JGI IMG server under proposal ID 503794.

