## Supplemental Methods and Figures for "Adaptive pangenomic remodeling in the *Azolla* cyanobiont amid a transient microbiome"

Once a fern strain had achieved a suitable biomass, we removed approximately 8 grams of plant material from the culture and rinsed it in sterile ultrapure water before removing the roots. Ferns were then agitated for 30s in an Erlenmeyer flask containing 0.2 mL of Triton X-100 in 200 mL of sterile ultrapure water. The contents of this flask were sieved through a flame-sterilized 1 mm steel mesh and then rinsed with sterile ultrapure water for two minutes until all Triton-X had been removed. This material was then divided among the 6 wells of a sterile tissue culture plate containing an enzymatic solution of Cellulase “Ozonuka” R-10 (1 gram), Macerozyme R-10 (0.5 g), Pectinase from *Aspergillus niger* (0.05 g), KH_2_PO_4_ (24 g), Dithiothreitol (0.077 g), and Polyvinylpyrrolidone (0.5 g) dissolved in 50 mL of 0.5M mannitol solution. The plate was placed into a vacuum chamber for tissue infiltration by the enzyme solution for 30-45 minutes at 20 kPa. The solution and plant material were then moved into a 50 mL centrifuge tube and shaken at 32° C at 80 rpm for 18 hours. The resulting macerated *Azolla* material was strained through clean 500 μm Nitex mesh into a sterile glass flask, and material remaining in the mesh was gently rinsed through the mesh with sterile 0.5M mannitol solution. This process flushes the separated leaf pockets into the flask, which, after being allowed to settle, were concentrated by decantation. The leaf pockets were then gently poured into a sterile petri dish and pipetted under a stereo microscope into a single 2 mL microtube containing phosphate buffer solution. This process was repeated for each *Azolla* strain.

For DNA extraction, leaf pockets collected from each strain were centrifuged and resuspended in 425 μL TE buffer, to which 10 μg/ml RNAse A and 5 mg/mL lysozyme were added. Following a 20 min incubation at 37° C, Proteinase K (100 μg/ml) and 50 μL 10% SDS were added and incubated at 50° C for 2 hours. Next, 250 μL phenol and 250 μL chloroform/isoamyl alcohol (24:1) were added to the tube, mixed gently, and centrifuged at 15000 × g for 2 min. The aqueous supernatant was removed for a second extraction. This step was repeated three times, after which 500 μL chloroform/isoamyl alcohol was added, gently mixed, and centrifuged again for 2 minutes. The aqueous layer was removed and the process was repeated. DNA was then precipitated in cold ethanol and resuspended in 50 μL TE buffer.

*Scanning electron microscopy*

Ferns were fixed in 2.5% glutaraldehyde for 1 hour at room temperature and then stored in the fridge at 4°C until further processed. The samples were washed thrice with water for 5 minutes each, post-fixed with 2% OsO_4_ then washed again six times in water. They were incubated in 25% and 50% DMSO for 30 minutes each. Samples were split open while submersed in liquid N_2_ with a pre-cooled knife by knocking with a hammer. Split leaves were then transferred to 50% DMSO to warm to room temperature and dehydrated stepwise in ETOH (60%, 70%, 80%, 90%, 100% 3x) for 5 min each step. Samples were then transferred to 50% T-butanol/EtOH for 5 min then to 100% T-butanol. They were frozen at -20°C and freeze-dried in a HITACHI ES-2030 freeze drier, mounted on stubs and sputter coated with gold, and imaged on a Jeol JSM-7900F Scanning Electron Microscope.

**SUPPLEMENTAL FIGURES**

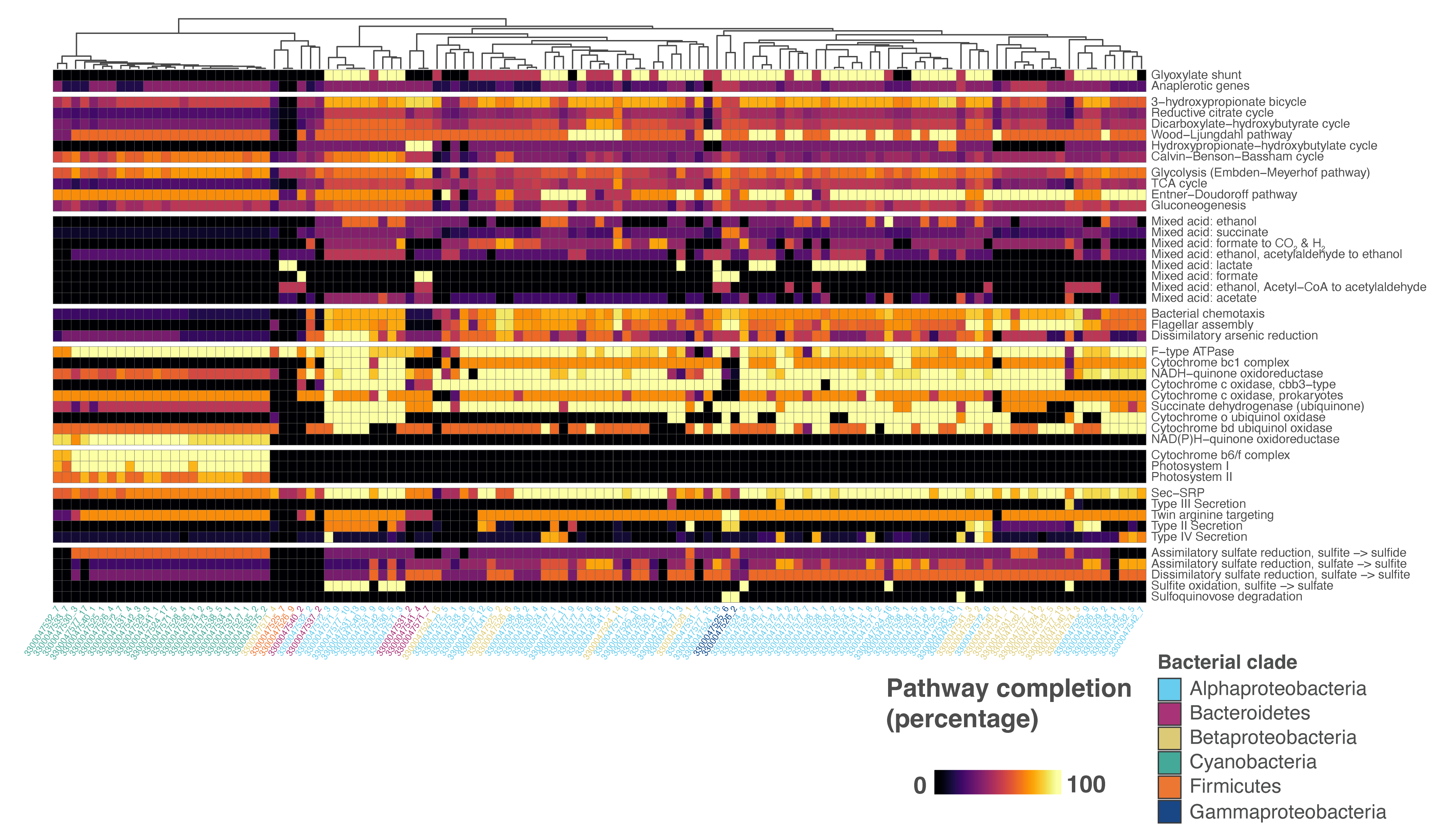

**Fig. S1**. — Heatmap of completeness of key metabolic and physiological pathways (rows) among all MAGs (columns) from the *Azolla* leaf pockets. MAGs are hierarchically clustered by functional pathway similarity following the dendrogram on top. Nitrogen transformation pathways are not pictured as they are already shown in figure 2, but also contribute to the clustering.

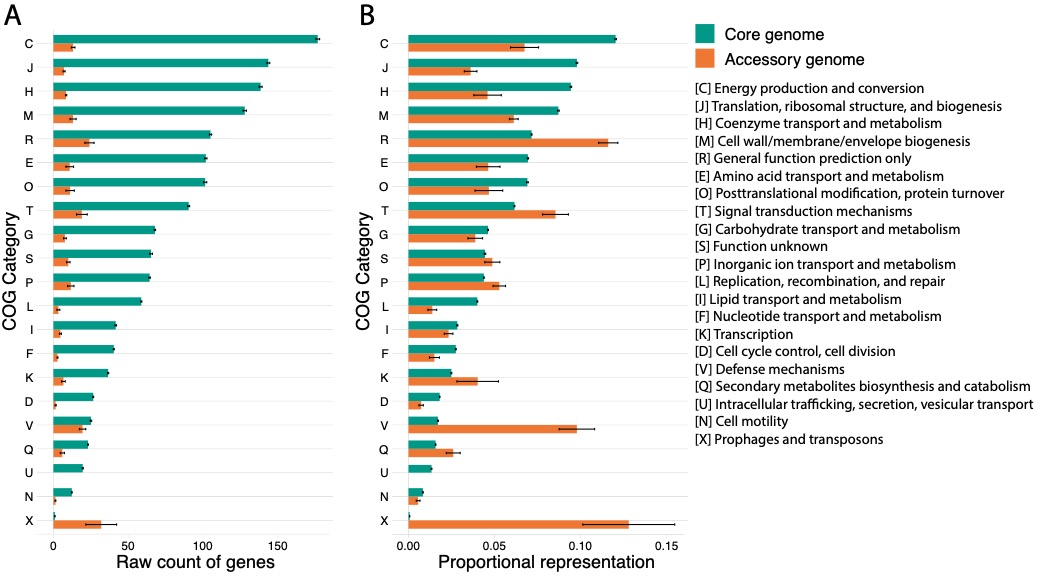

**Fig. S2**. — (A) Raw counts of orthologous gene clusters in *T. azollae* sorted into their respective COG categories and location on the *T. azollae* pangenome (see Fig. 3 for core and accessory definitions). (B) Proportional representation of a particular COG category relative to all others within the core or accessory genomes. Here, the accessory genome appears enriched in COG categories T, V, and X.

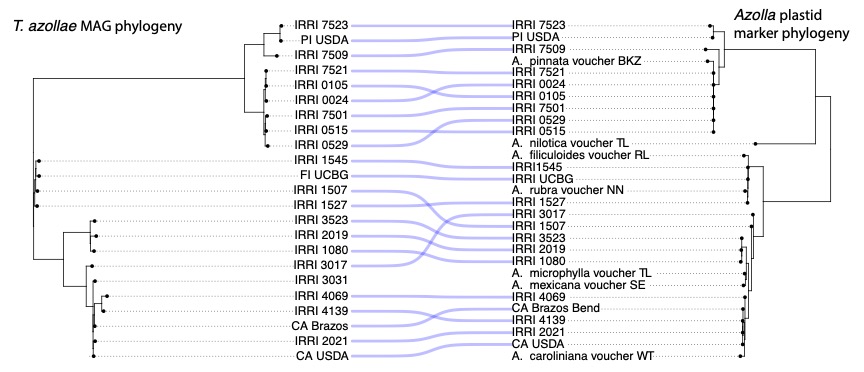

**Fig. S3**. — Cophylogeny of *T. azollae* MAG phylogeny and plastid marker phylogeny of the host fern *Azolla*. Voucher specimens are pulled from NCBI Genbank accessions and were not sequenced as part of the present study.
